# Assessment of Nutrient Composition and utilization of some popular wild Edible fruits of Kumaun Himalayan Region as Anti-oxidant agents

**DOI:** 10.1101/2023.06.21.545978

**Authors:** Neelaxi Pandey, Puja Ghosh, K. M Muhasina, Satpal Singh Bisht, Abhishek Jha

## Abstract

Food and Nutrition security is the main problem faced by developing and underdeveloped countries. Proper utilization of wild edible plants for nutrition security is a great choice for reducing the wastage of these powerful foods. The present investigation has been made to understand the nutritional and phytochemical parameter of selected underutilized fruits such as *Diospyros kaki, Pyrus pashia, Ficus semicordata, Diploknema butyracea, Pyracantha crenulata*, and *Rubus niveus*. The proximate analyses showed that *Diospyros kaki* is most promising fruit with 44.86± 0.4 mg/g carbohydrate, 9.29±0.80 mg/g protein, 81.31±0.4, 7% moisture and 1.1±0.11% little amount of ash content. Micronutrient iron was quantify highest in *Pyracantha crenulata* (3.52±0.24mg/100g) and zinc in *Rubus niveus*(8.13±0.05 mg/100g). Phytochemical screening was recorded in the ethanolic and aqueous extract, in which Phenolic and flavonoid content were highest in ethanolic extract of *Rubus niveus;* 64.05±0.13mg GAE/gm, 108.83±2.93mg QE/ g extract respectively and tannin content was highest in case of *Diospyros kaki* (79.94±0.40mg TAE/100g of extract). The free radical scavenging activity of fruits have been analyzed by DPPH, H_2_O_2_ and NO assay; it was observed that the ethanolic extract of *Rubus niveus* fruit is most promising with an IC_50_value 16.97μg/ml. In contrast, aqueous extract scored IC_50_value 28.86μg/ml. The lowest IC_50_ value was found in aqueous extract of *Diospyros kaki* i.e., 81.9 μg/ml in DPPH assay. The potential usage of these plants will help the people of these regions to combat nutrient deficiency diseases. Further development of research may help us to come out with powerful functional foods using these fruits.

**Graphical Abstract:** 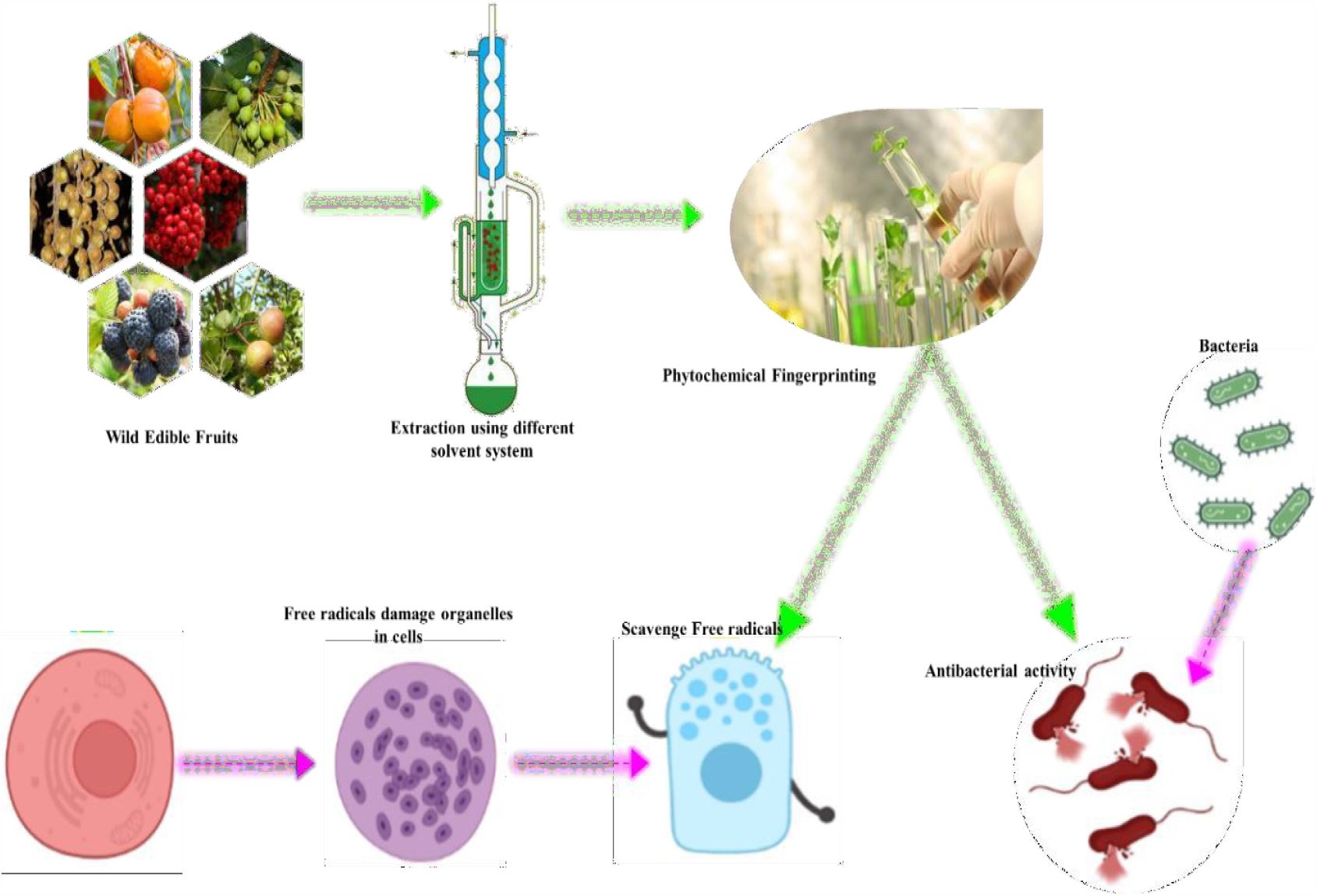

## 1. Introduction

Global food and nutrition security is a major challenging issue; it is believed that two billion people are suffering from malnutrition, which makes them more prone to diseases (Ritchie et al. 2018). Despite the fact that Africa and Asia are rich in plant diversity, including edible wild fruits, food security is a major concern in these regions. Plants have played a significant role in human culture since the time of the early hunter-gatherers and during various stages of adaptation. Many people around the world rely on wild species, particularly for food and medicine (Duguma 2020). There are thousands of underutilized edible plant species that are wild, semi-wild, or left out during domestication (Ray et al. 2020). The variety of edible wild plants found in India has the potential to revolutionize our food systems (Hunter and Fanzo, 2013; Powell et al. 2015). People frequently consume a wide range of uncultivated, wild plants as well as their parts, including flowers, fruits, green shoots, underground edible bits, and seeds.

For rural and semi-urban civilizations, wild edible fruits will improve the nutritional quality and offer an affordable source of nutrients. The main reasons for recommending these various diets are for optimum nutrition, sound health, and wellbeing. In rural and semi-urban areas around the world, both tribal groups and non-tribal populations have a long tradition of using wild edible plants for food and healing (Mahapatra et al. 2012).

During the 20th century, researchers were mainly engaged in studying the nutrient composition and ethno botanical perspective of plant and plant-based products. However, later on, advanced studies gained momentum due to the use of advanced tools and techniques, and investigations were initiated on pharmacological actions, food science, economic status, and microbiology of the plants and their products (Sardeshpande and Shackleton 2019). Fruits are nutrient-dense foods and offer vital components of a balanced diet. The term “nutrient-dense” refers to foods that are high in nutrients (such as potassium, magnesium, vitamin C, vitamin A, and fo<u>l</u>ate), dietary fiber, and other bioactive constituents. They are expected to have more positive effects on health while being rich in nutrition and low in calories (Slavin and Lloyd 2012). An extravagant consumption of nutrient-dense food is reported to have therapeutic activity for preventing and managing some chronic diseases, particularly cardiovascular diseases such as heart attack, stroke, diabetes, other degenerative, age-related diseases, and specific cancers. An increase in the consumption of fruits would be beneficial, especially for the elderly, who are more nutritional deficits than younger ones.

According to the Global Hunger Index report 2022, India ranks 101 out of 116 nations due to severe malnutrition. Three out of every ten children in India are stunted, and 43% of children under the age of 5 are underweight. Particularly in Uttarakhand, 26.6% are underweight, 33.5% are stunted, and 19.5% are wasted children, respectively. These malnutrition parameters are comparatively more in rural areas than urban areas (Rehan et al. 2020); food cost is the main barrier to consuming marketed foods, and that is why rural and tribal populations are unlikely to depend on wild fruits. Instead, traditionally gather fruits from the wild for their flavor, culturalpurposes, as food supplements, or to make up for food shortages. In addition, wild plants referred to as “famine food” or “hunger food,” may be used to address household food and economic security needs.

Uttarakhand is considered the floral basket of India, with 605 plants from 94 families. The Indian Himalayas flourished with 1748 commercially significant plants, comprising indigenous (31%) endemic (13.5%), and threatened (14%) species. The entire list of species from the Indian Himalayan region’s Red Data Book demonstrates the distinctive richness of such significant plants. Because of its challenging geology, harsh weather circumstances, and incredible flora, Uttarakhand is one of those locations suited for collecting wild edible fruits and plants. The people of this region rely on these plants as a great source of sustenance. Many wild fruits, including *Diospyros kaki, Pyrus pashia, Rubus niveus, Ficus semicordata, Pyracantha crenulata* and *Diploknema butyracea* have been exploited from the wild for centuries across the hills of Kumaun on account of its nutraceutical properties.

## 2. Methods

### Plant Material

According to the fruits’ seasonal availability, six wild edible fruits were collected from Nainital rainforest and Pithoragarh region of Uttarakhand, India. The fruits’ voucher specimen numbers are: *Diospyros kaki* (134), *Diploknema butyracea* (135), *Ficus semicordata* (136), *Pyracantha crenulata* (137), *Rubus niveus* (138), and *Pyrus pashia* (139). (B.S.I.Certificate).

### Preparation of the raw material

All collected fruit samples were thoroughly washed with water, followed by mechanical segregation and then oven-dried at 38^0^C with varied time durations. The dried fruit material was further ground to powder and stored in zip-lock poly bags to avoid possible moisture contamination. For fruits of *Diploknema butyracea* and *Dysporous kaki* due to their liquefying and oily consistency, oven drying was not satisfactory; therefore, these fruits were further chopped into small pieces and stored in zip-lock poly bags under moisture-free conditions.

Finely chopped fruit content was extracted with the help of the Soxhlet apparatus in increasing solvent polarity (ethanol and water). Using a rotary evaporator (IKA, Germany), each extract was concentrated under vacuum. All extracts were kept at 4°C.

### Estimation of total phenolic content

The total phenolic content was ascertained using the Folin-Ciocaltaeu assay (Chandran and Indira 2016). Study’s standard was gallic acid At 760 nm, the absorbance was measured in reference to a typical blank solution made up of all the reagents using 1 ml of plant extract from various concentrations that had been diluted ten times with 5 ml of Folin-Ciocaltaeu reagent, incubated for 2 minutes, and then added 5 ml of 7.5% sodium carbonate solution. Gallic acid milligrams per gram of dried fruit material was used to measure the phenolic content.

### Estimation of total flavonoid content

Quercetin was used as a standard to assess the total flavonoid concentration (Chandran and Indira 2016).Take a volumetric flask, add 300μl of 5% sodium nitrite with 100 μl of different quercetin concentrations, and 4 ml of H_2_O. After five minutes, a 300μl solution of 10% aluminium chloride was added, six minutes later, 2 ml of 1M sodium hydroxide. The solution was filled with distilled water to a total of 10ml. At 510 nm, the absorbance of the solution was measured in reference to a blank solution, contained all reagents except aluminium chloride.

### Estimation of total tannin content

The Folin-Ciocalteu method is used to estimate total tannins (Chandran and Indira 2016). The standard was tannic acid. In a volumetric flask containing 7.5 ml of distilled water, 500μl of Folin-Ciocaltaeu reagent, and 1 ml of diluted 35% sodium carbonate solution, 100μl of plant extract and various standard quantities were added. The solution’s absorbance was measured at 700 nm in comparison to a control after the components had been mixed and held at room temperature for 30 minutes. The amount of tannin was expressed as mg tannin acid equivalent per gram (TAE/g).

### Estimation of Total Moisture Content

A fruit sample’s moisture content, frequently referred as water content, indicates how much water is actually present in the sample. According to the traditional definition, moisture content is the percentage-based relationship between the mass of water and mass of solids in a sample.

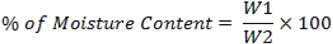

W1=Weight of dried fruit sample

W2= Weight of fresh fruit sample

Firstly weigh 10g of fresh fruit sample then dry in an oven until constant weight is achieved (Park, 1996).

### Ash Values

Ash is the indication for the byproduct left over/after the burning of the raw drug. The residue is an indication of the inorganic salts that are naturally present in the fruit and stick to it. As a result, determining the ash value provides the foundation for determining the authenticity and cleanliness of a sample.

In a tarred silica crucible, 10 g of oven-dried powdered fruit sample was used and it was incinerated in the muffle furnace for one to two hours. All organic material burns at this temperature except minerals; the crucible was taken out and cooled. To get at a constant value the process was repeated after another weigh. Finally, In relation to the air-dried fruit sample, the percentage of total ash value was calculated (Park 1996).

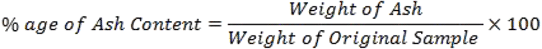

### Total Protein Estimation

The Brad-fords assay was used to quantify proteins (Bradford, 1976). Fruits were collected and cleaned under running water for the quantitative protein analysis. First, a sample of 5 g of fruit was taken, and using a mortar and pestle, the sample was extracted in phosphate buffer. The fruit sample was mashed into a smooth, thick paste. The smooth paste was then collected in a 1.5 ml microfuge tube and centrifuged for 30 minutes at 4000 round per minute (rpm). The upper layer supernatant was collected and kept at 4°C for further use after centrifugation. It was dyed with Coomassie brilliant blue G-250, which mostly binds to the amino acid residues in proteins (arginine, histidine, lysine, tyrosine, tryptophan, and phenylalanine). The dye’s absorbance in the ultraviolet (UV) spectrometer shifts from 465 to 595 nm as a result of this binding to protein. As a result, the optical density was assessed using U.V. spectroscopy at 595 nm to determine the sample’s protein content.

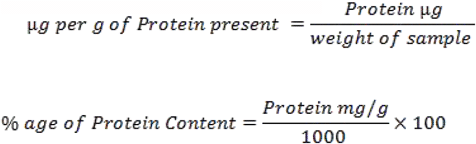

### Estimation of Carbohydrates

Fruit samples were first hydrolyzed into simple sugars using diluted hydrogen chloride to estimate the amount of carbohydrates. Then, glucose was dehydrated into hydroxymethylfurfural by dehydrating it in a hot acidic medium (HMF). At 620 nm, it combines with anthrone to generate a green colour complex; this green complex ensures the presence of carbohydrates.

From the 200 g/ml stock solution, varied amounts of the glucose solution were pipetted into several test tubes. Tubes 2 through 6 and tubes 7 through 12 were set up for a standard curve, and tube 1 served as a blank. Test samples were used in tubes 7 through 12. Each test tube received five ml of the supplied anthrone reagent, which was completely mixed in by vortexing. All test tubes were heated to 90 degrees Celsius for 17 minutes under cover after being given time to cool. An optical density (O.D.) measurement at 620 nm was taken after it had reached room temperature against a blank. The computations used a standard curve that was created.

### Iron and Zinc Estimation by Atomic absorption spectroscopy (AAS)

The iron and zinc content of six edible wild fruits evaluated by AAS using the iron and zinc standard (Sigma Aldrich). In reference to known concentrations of iron and zinc (1 g/ml, 2 g/ml, and 10 g/ml from the stock solution of 1000 g/ml), a standard curve was developed (Beaty and Kerber 1993).

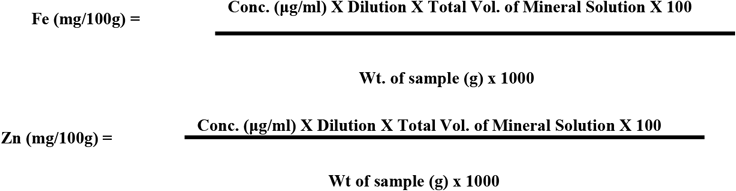

### Antioxidant Assay

Through the use of the DPPH Assay, Nitric Oxide Assay, and Hydrogen Peroxide Assay, free radical scavenging activity was examined.

### 2, 2-Diphenyl-1-Picrylhydrazyl (DPPH) Activity

By using the DPPH assay, free radical scavenging activity was calculated (Williams *et al*., 1995). Ascorbic acid used as the reference. 800 ml of DPPH solution (using methanol, DPPH molarity was 90 μM) was added to 200 μl

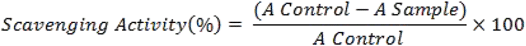

of fruit extract in volumetric flasks at different concentrations. The solution was allowed to stand at room temperature for 30 minutes before being measured at 515 nm in reference to a blank. Methanol and DPPH solution were typical components of a blank solution. The following formula was used to calculate scavenging activity. To determine antioxidant capacity, the IC_50_ value was calculated using linear regression analysis.

#### 2.10.1. Nitric Oxide Assay

In an aqueous sodium nitroprusside solution with a pH of 7.2, nitric oxide spontaneously forms. When nitric oxide reacts with free oxygen, nitrate and nitrite are produced as a stable byproduct. The Griess reagent can be used to measure these stable ions. Cellular harm could occur if nitric oxide production is excessive.

The assay was carried out largely in accordance with the procedure described by (Parul et al. 2013). Extracts were obtained in various test tubes with different contents, such as 20, 40, 60, 80, 100, and 120 μl, and ascorbic acid was employed as the standard. Following the addition of sodium nitroprusside (5 mM) in phosphate buffered saline, the volume was increased to 2 ml with DMSO. Each tube received 2 ml of Griess reagent (1% sulphanilamide, 0.1% naphthyl ethylenediamine dichloride, and 3% phosphoric acid) after two hours of incubation at 30 °C, and the absorbance at 546 nm was immediately noted.

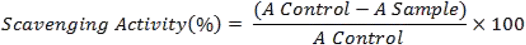

#### 2.10.2. Hydrogen Peroxide Assay

The method described by Sasikumar and Kalaisezhiyen (2014) was used to test the fruit extracts’ capacity to scavenge hydrogen peroxide, with a few minor modifications. Hydrogen peroxide solution (43 mM) was made in PBS buffer (0.1 M, pH 7.4). The sample was added to the hydrogen peroxide solution at various doses (10-100 mg/ml) (0.6 ml, 43 mM). After 10 minutes, the reaction mixture’s absorbance value at 230 nm was recorded in reference to a phosphate buffered blank solution devoid of hydrogen peroxide. Ascorbic acid served as the benchmark.

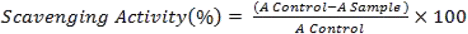

As mentioned previously, the free radical scavenging activity was calculated using the percentage of inhibition.

## 1. Results & Discussion

The characteristics of food, food sources, and other substances that are experienced by the senses of the human body, such as taste, sight, smell, touch, colour and size are referred to as organoleptic properties. Six edible wild fruits that were the subject of this study’s organoleptic analysis are listed in Table 3.1. Understanding or determining if food or pharmaceutical goods can transfer tastes or odours to the materials and components they are packed in requires organoleptic studies.

### 3.1. Total Phenolic Content

Polyphenol content has an aromatic ring structure with a hydrogen donating group (Huaman-*Castilla et al*., 2020). A fruit’s quality and nutritional worth are generally influenced by its phenolic components, which also affect the fruit’s colour, flavour, and taste (Vaya and Aviram 1997). Additionally, phenolic chemicals are in charge of a plant’s colour, development, reproduction, and defence mechanisms, which guard against cellular harm from microbes, insects, and herbivores (Lattanzio *et al*., 2006). The primary cause of phenolics’ antioxidant action is their redox potential, which lowers the concentrations of harmful and foreign substances, hydrogen donors, singlet oxygen quenchers, and metal chelators (Nagarajan *et al*., 2020). The Folin-Ciocalteu method, as described by Chandran and Indira (2016) with some minor modifications, was used to determine the total phenolic content; the results are reported as Gallic acid equivalents in milligrammes per gram of extract.

Figure 3.1 provides an analysis of the total phenolic content in various fruit extracts. The results of the current investigation showed that the ethanolic extract had a larger phenolic content than the aqueous extract. Per gram of the extract, the total phenolic content ranged from 13.51± 0.48 to 64.05±0.13 of gallic acid equivalent. Out of the twelve extracts examined in the current experiment, the total polyphenol content of *Rubus niveus* fruit in the ethanolic extract was determined to be at its highest (64.05% + 0.13). The aqueous extract of *Diploknema butyracea* had the least amount of polyphenols (13.510.48). With ethanol, various extracts’ phenolic concentration reduced in the following ways: RNE> DKE> PCE> FSE> PPE> DBE and water extract RNW> PCW> FSW> DKW> PPW>DBW, respectively.

**Figure 3.1:**
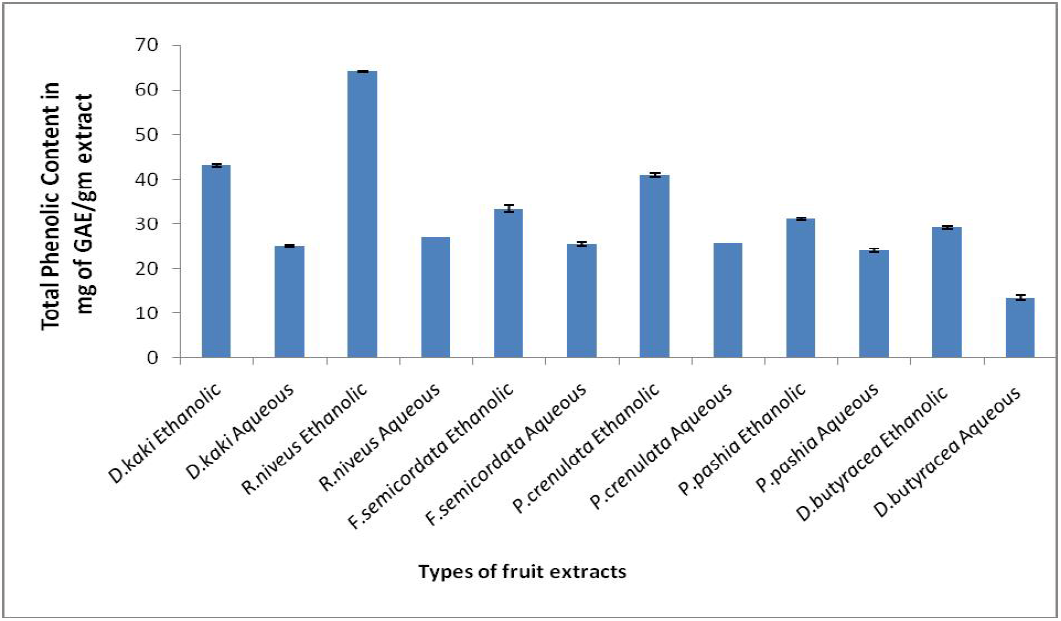
Total Polyphenolic Content of Six Wild Edible Fruit Extracts.

### 3.2. Total Flavonoid Content

Flavonoids are the most abundant secondary metabolites of fruits, total flavonoid content of extracts was evaluated following Chandran and Indira (2016) with minor modifications.

The highest flavonoid content was found in the ethanolic section of *Rubus niveus*

*i*.*e*. 108.83±2.93 mg of QE/g of extract and the lowest was reported in the case of the aqueous extract of *Diploknema butyracea* i.e. 4.44±1.77 (fig 3.2).

However, the total flavonoid content in the present study ranged from 4.44±1.77 to 108.83±2.93 of quercetin equivalent per gram of extract. The ethanolic and aqueous extracts of *Diospyros kaki* and *Pyrus pashia* recorded 40.83±1.79, 63.11±0.60, 4. 52±0.47 and 12.16±0.20 mg of quercetin equivalent /g of extract respectively. The flavonoid content in the ethanolic extract of *Rubus niveus* was 108.83±2.93 mg of quercetin equivalent /g of extract. All these observations indicate that these WAFs are the candidate source of flavonoids in the hills of the Kumaun region. The total flavonoid content of different extracts decreased in the following manner with ethanol i.e. RNE> FSE> PPE> PCE> DKE> DBE> and water extracts RNW> FSW> PPW> PCW> DKW>DBW respectively.

**Figure 3.2:**
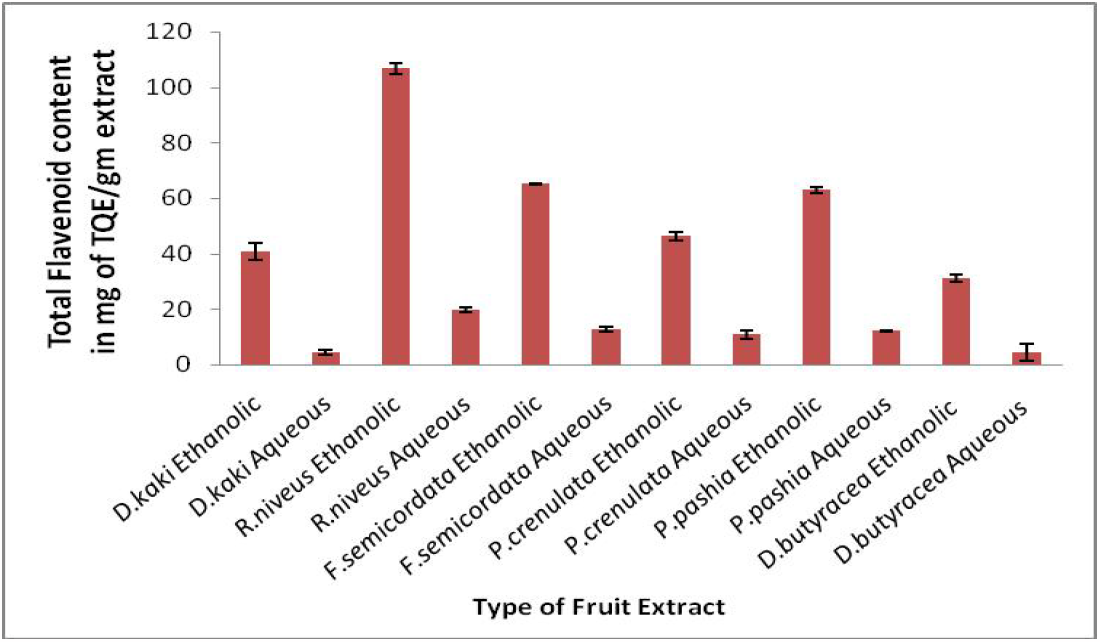
Total Flavonoid Content of Six Wild Edible Fruit Extracts.

### 3.3. Total Tannin Content

Tannins from a group of phenolic compounds result from flavonoid units’ polymerization. These are also antifeedants with antidigestive properties, which acquire protein binding in the diet and make them inedible (Masette *et al*., 2015). Total tannin content estimated during the present investigation from selected WEFs revealed that the highest tannin content is present in ethanolic extract of *Diospyros kaki*, followed by *Pyrus pashia* and *Rubus niveus* Fruit.

**Figure 3.3:**
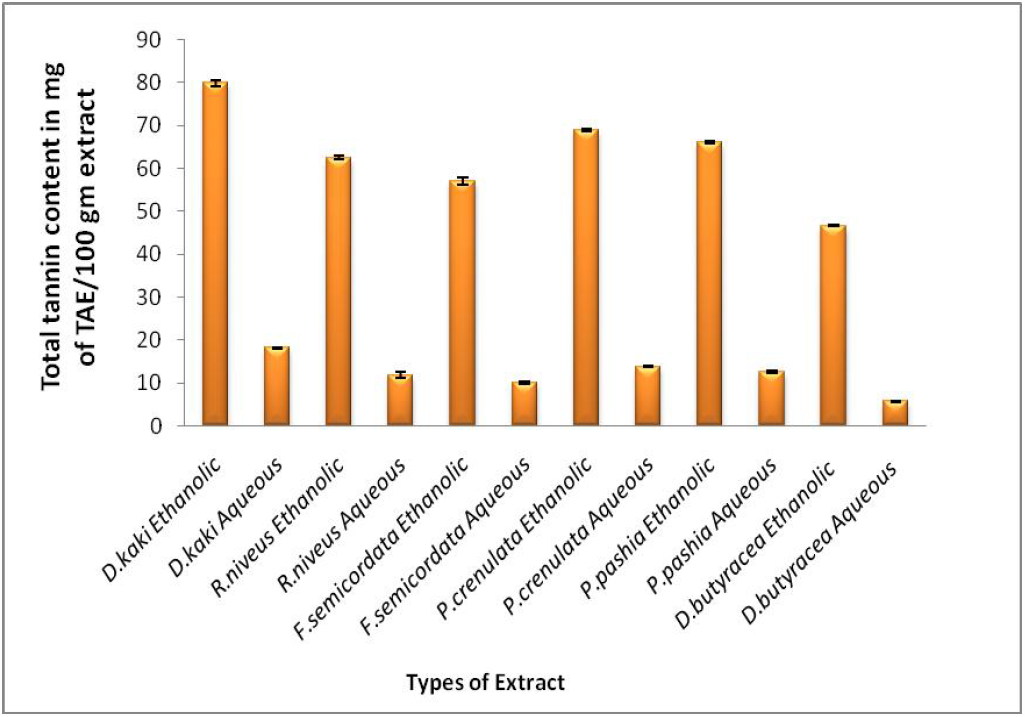
Total Tannin content Acid Content of Six Wild Edible Fruit Extracts.

### 3.4. Nutritional Analysis of Fruits

Fruits are a key component of a balanced diet and provide numerous health advantages when consumed regularly. In the current experiment, a number of nutritional factors were examined in order to determine the nutritional value of the six edible wild fruits. All fruits’ percent moisture content values ranged between 40.81± 0.26% (Diploknema butyracea) and 81.31±0.47% (*Diospyros kaki)*. The higher moisture content in these wild edible fruits could be related to their habitat. Additionally, there is a clear correlation between fruit ash content and mineral content, *Diploknema butyracea* contains 3.37±0.30% of ash, whereas *Diospyros kaki* fruit consists of 1.1±0.11%, the ash content of remaining fruit samples observed between 3.37±0.30% to 1.1±0.11%. At all levels of metabolic activity, energy is produced by the macromolecules, proteins, and carbs that make up a large component of the food. The *Ficus semicordata* reported comparatively higher protein content (7.24±0.35 mg/g) with moderate (27.75±1.08 mg/g) carbohydrate value. However, *Diospyros kaki* gave lower protein i.e. 9.29±0.80 mg/g with high carbohydrates (44.86±0.49 mg/g). The results regarding the proximate composition of selected wild edible fruits are presented in Table 2.

The higher amount of carbohydrate in *Diospyros kaki* makes it an excellent instant food and energy source for the people; similarly, a low value of carbohydrate is reported in the case of *Diploknema butyracea* making it an acceptable food source for people having metabolic disorders like diabetes type II. The proximate analysis revealed that *Pyracantha crenulata* can be used as protein-rich WEFs with high digestibility and fiber content. The analysis showed that all these fruits have their unique composition of macro and micronutrients *viz. Rubus niveus* is the very common wild blackberry of the Himalayan mountain region, a rich source of Zn. That makes it a very promising fruit responsible for immunity and immunomodulator.

### 3.5. Antioxidant Activity

Reactive free radicals and its detrimental effect in foodstuff and lipid rancidity, cellular damage, and illnesses are currently the subject of attention. Numerous other investigations have confirmed that free radicals play a role in the spread of lipid oxidation and that many reactive free radical species are produced throughout a variety of metabolic activities. Therefore, when evaluating the activity of radical scavenging, relatively stable radicals as DPPH• are frequently used.

The results of using the standard approach with a minor changes. IC50 value 16.97 g/ml, the alcoholic extract of Rubus niveus berries is the most promising of the fruit extracts to scavenging free radicals while aqueous extract in comparison, has an IC50 value of 28.86 g/ml. The lowest IC50 value was found in an aqueous extract of *Diospyros kaki* i.e., 81.9 μg/ml. The total antioxidant potential of different extracts observed as mentioned below RNE(16.97)>PPE(18.58)>PCE(18.75)>FSE(22.02)>RNW(28.86)>FSW(32.44)>PCW(37.53)>PPW(46.46)>DBE(4 7.27)>DKE(50.51)>DBW(69.8)>DKW(81.9).

Calculating the decreasing absorbance at 517 nm caused by the scavenging activity of stable DPPH free radicals was done to determine the free radical scavenging activity. The test samples may have the ability to scavenge free radicals, according to the results of the positive DPPH tests.

The presence of certain chemicals and structural elements, such as the quantity of phenolic hydroxyl or methoxyl groups, flavone hydroxyl groups, keto groups, free carboxylic groups, and other structural characteristics, affect an extract’s antioxidant activity.

While comparing all three assays to determine the antioxidant activity, sizable variations were observed in IC50 value for Diospyros kaki extracts as it yielded almost similar results in three assays. However, a significant difference was observed in ethanolic extract of *Rubus niveus* IC50 value 16.97 (DPPH assay), 50.56 (H2O2 assay) and 47.27 (NO assay). The IC50 value for different WEFS in various solvents is tabulated in fig 3.1.

All the fruit extracts showed their antioxidant potential (I.C. 50 value), ranging from 59.56 to 89.68. Generally, ethanol extracts show high potency against Hydroxyl free radicals than aqueous extracts. A maximum IC50 value was offered by aqueous extract of *Diploknema butyrecea* (89.68) and *Pyrus pashia* (86.67) and the lowest value was reported in ethanolic extract of *Rubus niveus* i.e. 59.56.

The bioregulatory molecule nitric oxide is essential for many physiological activities, including the transmission of brain signals, immunological response, control of vasodilation, and blood pressure. It functions as an effector in a variety of processes, including neuronal messenger, vasodilation, and antibacterial and anticancer actions. Upon stimulation with cytokines and bacterial lipopolysaccharide, NOS are generated in a range of cell types and they consistently produce large quantities of nitric oxide.

One of the most popular and effective tests for neutralizing free radicals is nitric oxide. On the basis of current investigation, its values ranged from 47.27 to 164.9. In general, ethanol extracts shows effective results over aqueous extracts in terms of their ability to scavenge nitric oxide. Maximum scavenging activity was provided by aqueous extracts of *Pyracantha crenulata* (164.9) and *Rubus niveus* (47.27). The hyperactivity is stabilized by these samples’ interactions with nitric oxide free radicals.

**Fig. 3.4.**
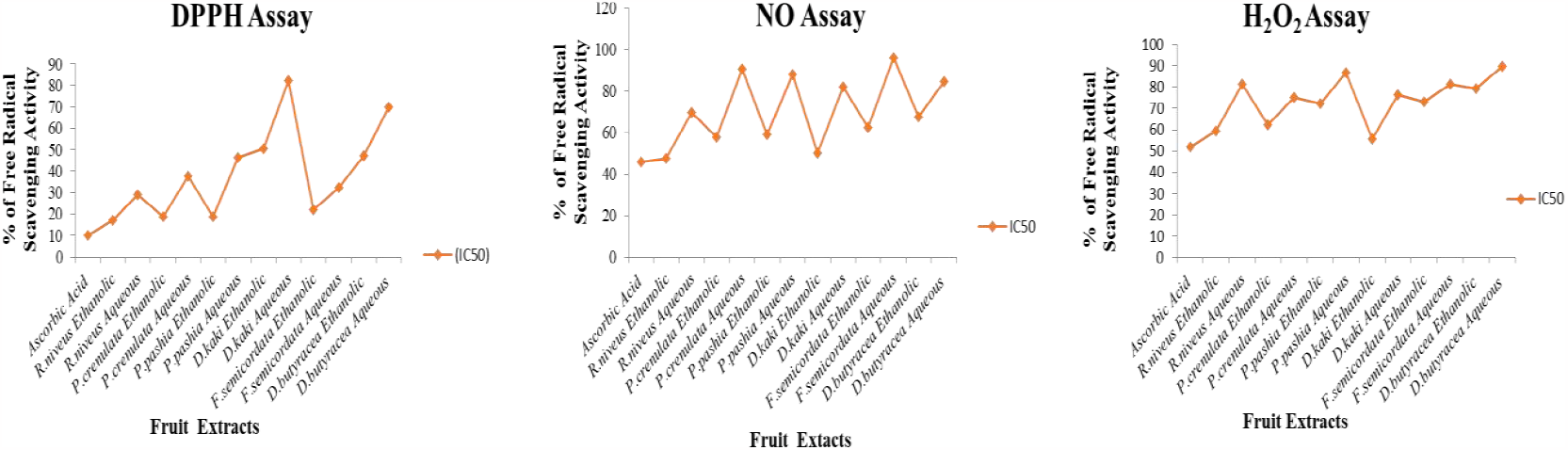
Free Radical Scavenging Activity of Six Wild Edible Fruit Extracts.

## 3. Conclusion

Fruits are an essential constituent of a balanced human diet to maintaining a healthy life. The health-promoting nutraceutical effects are associated with their intakes, primarily due to secondary metabolites. Wild edible fruits have been established as a suitable source to deal with many chronic diseases, such as diabetes, obesity, cardiovascular diseases, and lung, breast, skin cancers. The six underutilized wild edible fruits are bestowed with potent antioxidant and antibacterial activity. The bioactive constituents present in these fruits, especially phenolics and flavonoids, help reduce free radical production and antibacterial activity. These fruits have more therapeutic potential than marketed fruits and are cost-effective. This study may help the researchers and food processing industries working in functional and fortified foods.

## 4. Acknowledgement

I would like to thank my all the authors for contributing in manuscript and Kumaun University for providing the research facilty.

## 5. Conflicts of Interest

The authors declare that there is no conflicts of interest exist.

## 6. Funding

I am thankful to Kumaun University for providing the research Facility

## Abbreviations

GAE: Gallic acid equivalent
QE: Quercetin equivalent
TAE: Tannic acid equivalent
IC_50_: Half maximal inhibitory concentration
DPPH: 2,2-diphenyl-1-picrylhydrazyl
H_2_O_2_: Hydrogen peroxide
NO: Nitric oxide

## Notes

### Competing Interest Statement

The authors have declared no competing interest.

